# Plant phenology supports the multi-emergence hypothesis for Ebola spillover events

**DOI:** 10.1101/158568

**Authors:** Katharina C. Wollenberg Valero, Raphael D. Isokpehi, Noah E. Douglas, Seenith Sivasundaram, Brianna Johnson, Kiara Wootson, Ayana McGill

## Abstract

Ebola virus disease outbreaks in animals (including humans and great apes) start with sporadic host switches from unknown reservoir species. The factors leading to such spillover events are little explored. Filoviridae viruses have a wide range of natural hosts and are unstable once outside hosts. Spillover events, which involve the physical transfer of viral particles across species, could therefore be directly promoted by conditions of host ecology and environment. In this report we outline a proof of concept that temporal fluctuations of a set of ecological and environmental variables describing the dynamics of the host ecosystem are able to predict such events of Ebola virus spillover to humans and animals. We compiled a dataset of climate and plant phenology variables and Ebola virus disease spillovers in humans and animals. We identified critical biotic and abiotic conditions for spillovers via multiple regression and neural networks based time series regression. Phenology variables proved to be overall better predictors than climate variables. African phenology variables are not yet available as a comprehensive online resource. Given the likely importance of phenology for forecasting the likelihood of future Ebola spillover events, our results highlight the need for cost-effective transect surveys to supply phenology data for predictive modelling efforts.

## Introduction and purpose

Ebola Virus Disease can emerge simultaneously at geographically distant locations throughout the West African tropical rainforest belt biome. For example, the 1996 wave of outbreaks in both animals and humans occurred within just a few months in remote forest locations in Gabon and the Democratic Republic of Congo (Georges et al. 1999; Milleliri et al. 2004; Yamagiwa and Basabose 2006; Lahm et al. 2007). This fact evoked migrating species such as bats as possible natural reservoirs of the virus (Leroy et al. 2004; Leroy et al. 2009). However, most emergences of Ebola virus disease have not been linked to direct human exposure to bats (Leendertz et al. 2016), and global predictions do not identify the African tropical rainforest belt biome as hotspots for viral spillovers from bats (Olival et al. 2017). Since current evidence for migrating bats being the primary natural reservoir of Ebola virus is inconclusive, alternative explanations for the sporadic emergence of Ebola virus disease in different places at the same time should be explored. Phylogenetic distance between susceptible potential host species has recently been used to predict additional susceptible taxa (Weber et al. 2017; Olival et al. 2017). Mononegavirales, the supergroup containing Filoviridae such as the Ebola virus group, include a broad spectrum of natural reservoirs, including many mammals and other tetrapods (Taylor et al. 2010), so that no potential reservoir species can be *a priori* excluded.

Simultaneous infections of humans and animals in the Gabon-Congo border region between 2001 and 2005 originated from viral strains with different amino acid sequences of the viral glycoprotein (GP), indicating that these originated from at least five independent, simultaneous spillover events (Leroy et al. 2004; Wittmann et al. 2007). Based on this evidence, and the fact that viruses cannot exist outside of their natural host organisms, previous studies highlighted possible ecological and environmental (“eco-environmental”) dimensions to Ebola virus spillover events.(Wittmann et al. 2007) hypothesized that “particular, but unknown environmental conditions” may have caused the multiple spillover events from the unknown host to the human and animal populations recorded to be affected (Peterson et al. 2004). Leroy and Gonzalez (2012) also called this the “multi-emergence” hypothesis, where multiple, episodic, and simultaneous outbreaks are caused by ecological or environmental conditions.

An environmental niche model has been generated for the Ebola virus based on environmental parameters in its localities of emergence, showing the African tropical rainforest biome as clearly defined region with highest probability of occurrence (Pigott et al. 2014). Besides seasonal fluctuations, and despite annual bat migration and high rates of human consumption of bushmeat, Ebola virus disease does not emerge annually. The 1990s Ebola outbreaks were related to short-term changes in regional precipitation and Normalized Difference Vegetation Index (NDVI) (Tucker et al. 2002). Evapotranspiration and Enhanced Vegetation Index are predictive of the spatial environmental niche of Ebola virus (i.e., being located in the African tropical rainforest biome), and monthly rainfall and rainfall anomaly can predict Ebola virus spillovers (Pinzon et al. 2004; Schmidt et al. 2017).

Intraspecific or interspecific competition for resources such as food, or environmental contamination within the virus-harbouring ecosystem can serve as a hypothetical functional link between ecological and environmental factors and spillover events from the unknown natural reservoir to other species. (Rouquet et al. 2005) hypothesized, that direct competition for resources between susceptible species in periods of food scarcity leads to increased inter-species transmission (Rouquet et al. 2005), and the annual fruit bat migration into the Democratic Republic of the Congo between April and May could be related to human disease emergence events in June (Leroy et al. 2009). A new mechanistic hypothesis invokes “viral rain”, the shedding of infectious viral particles from the natural reservoir into the environment, such as bats spreading Hendra viruses around the trees they roost on (McFarlane et al. 2011).

Current efforts to identify Ebola outbreaks exclusively rely on monitoring on the level of animal species commonly hunted as bushmeat, and first incidences in humans (Rouquet et al. 2005). This practice is costly and associated with complicated logistics. Designing novel methods for forecasting the likelihood of future Ebola virus disease spillover events before the human-to-human transmission stage is reached, would be a cost-effective method of upstream mitigation (Schar and Daszak 2014).

In this article, we evaluate support for the multi-emergence hypothesis from ecological and environmental data sourced from the zoonotic niche of Ebola virus (Pigott et al. 2014), as being predictive for Ebola virus disease spillover events. Our working hypothesis is, that local climate and phenology variables (greening, flowering, and fruiting) will be better predictors for Ebola virus disease spillovers than remotely sensed climatic variables currently used for predictive modelling.

## Methods

Data on the occurrence of Ebola virus spillover events were obtained from published reports (for localities see Figure 1a). Therefore, the true number of spillover events (especially in animals) might be higher, and we have to consider the dataset we compiled as a sample of this true number. Data for spillover events involving humans was obtained from the website of the United States Centers for Disease Control and Prevention (cdc.gov, accessed 2014-2017). As the CDC website only lists reports for human outbreaks, we searched online literature resources for information on other species’ Ebola infection records to obtain data on outbreaks in non-human animals (International Commission 1978; Heymann et al. 1980; Le Guenno et al. 1995; Muyembe and Kipasa 1995; Amblard et al. 1997; Formenty et al. 1999; Georges et al. 1999; Khan et al. 1999; Okware et al. 2002; Leroy et al. 2004; Lamunu et al. 2004; Milleliri et al. 2004; Rouquet et al. 2005; Pourrut et al. 2005; Lahm et al. 2007; Wittmann et al. 2007; Onyango et al. 2007; Towner et al. 2008; Leroy et al. 2009; Wamala et al. 2010; MacNeil et al. 2010; Nkoghe et al. 2011; Shoemaker et al. 2012; Bausch and Schwarz 2014; Baize et al. 2014). Wittmann et al. (2007) and Leroy et al. (2009) reported on molecular screens for Ebola virus strains for the 2001-2005 outbreaks, so we counted each confirmed independent epidemic chain as independent event. In total, our dataset includes 57 human and other animal spill-over events between 1976 and 2017.

**Figure 1.**
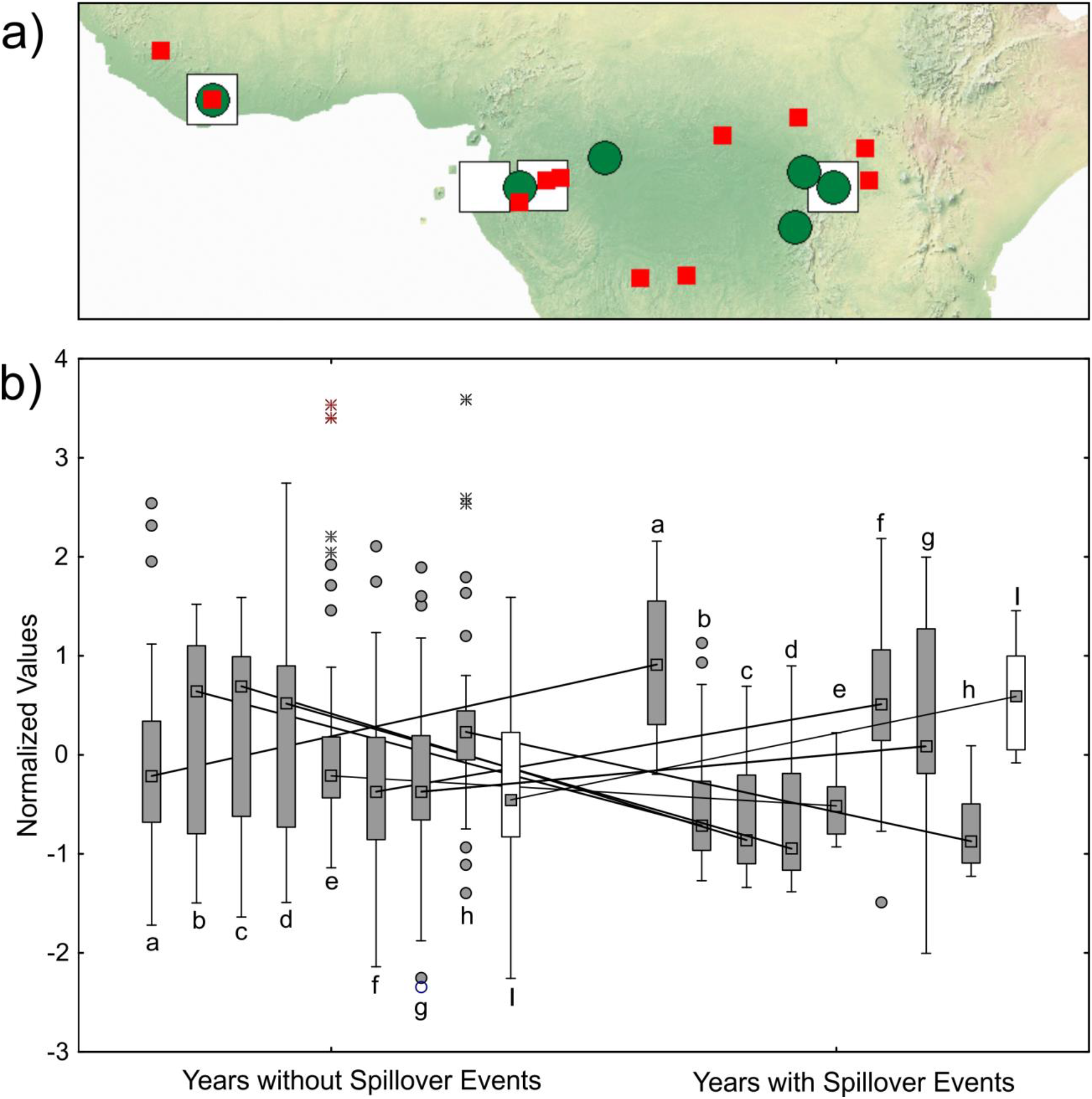
a) Map of localities of Ebola virus disease outbreaks (where exact coordinates could be obtained, small squares), climate data (large squares), and phenology data (circles). b) Box plots for climate and phenology variables with significant inter-annual variation (See Table 1 for statistical results) between years with recorded Ebola human + animal spillover events and years without known Ebola Virus spillover events spanning the period of 1953-2016. Boxes represent 25-75% interval, whiskers represent non-outlier range, circles represent outliers, and stars represent extreme values. Lines connect medians of the same variables between years with spillover events, and years without spillover events. Climate variables are shown in filled boxes, phenology variable is shown in open boxes. a) 10 Year average rainfall in Kibale National Park b) Number of days with ≥0.254 mm rainfall per month (DP01) c) Number of days with ≥12.7 mm rainfall per month (DP05), d) Number of days with ≥25.4 mm rainfall per month (DP10), e) highest daily total of precipitation per month in mm (EMXP), f) Lowest daily minimum temperature for the month in °C (EMNT), g) monthly mean minimum temperature in °C (MMNT), h) total precipitation, in mm/10, for the month (TPCP), i) Second Independent Component of Normalized Difference Vegetation Index (IC2 NDVI) anomaly between July and December.

Climate data was obtained from the Global Historical Climatology Network (GHCN, NOAA accessed 2015-2016, Figure 1a). Data was obtained for monthly summaries between 1953-2015 for the weather station GHCND-GB000004556 in Makokou, Gabon (Lat 0.578004°, Long 12.889751°), and for the years 2016-2017 for the location FIPS:GB 9 (Libreville, Gabon, Lat 0.457088°, Long 9.409372°). We obtained data for the following variables: DT90, Number of days with maximum temperature ≥32.2°C; DP01, Number of days with ≥0.254 mm rainfall per month; DP05, Number of days with ≥12.7 mm rainfall per month; DP10, Number of days with ≥25.4 mm rainfall per month; CLDD, cooling degree days, computed when daily average temperature is more than 18.3°C; CDD, days per month with mean daily temperature −18.3°C; EMNT, Lowest daily minimum temperature for the month in °C; EMXP, highest daily total of precipitation per month in mm; EMXT, highest daily maximum temperature per month in °C; MMNT, monthly mean minimum temperature in °C; MMXT, monthly mean maximum temperature in °C; MNTM, monthly mean temperature in °C; TPCP total precipitation, in mm/10, for the month. From this monthly dataset, we calculated arithmetic means for each year, as well for each month over the 1953-2017 period. We added additional climate data from other published sources, that were computationally collected from published graphs using the WebPlotDigitizer software (V 3.12) (Rohatgi 2016). Average annual values and for average rainfall (1941-2000), and average monthly minimal and maximal temperatures (1976-2001) for Kibale National Park, Uganda (Lat 0.486621°, Long 30.390715°) were obtained from Chapman et al. (Chapman et al. 2005). From Polansky and Boesch (Polansky and Boesch 2013), we extracted monthly inter-annual linear trends of rainfall, average monthly rainfall in Central Africa (in mm), between 1979-2010 in Tai National Park, Ivory Coast (Lat 5.690020°, Long - 6.939547°). We also extracted temperature for African humid tropics in °C (Fisher et al. 2013), as well as the averaged normalized departure (Apr-Oct rainfall departure) between 1979-2010 (Boyd et al. 2013).

Currently, no comprehensive database exists for African plant phenology and vegetation data (Adole et al. 2016). The African tropical rainforest belt, which has high occurrence probability for Ebola virus disease emergence (Pigott et al. 2014), is characterized by a regional, seasonal pattern of greening, fruiting and flowering linked to the cyclical West African Monsoon (Cornforth 2013). We used WebPlotDigitizer (Rohatgi 2016) to collect quantitative plant phenology datasets in published reports from the area of high occurrence probability for the Ebola virus (Figure 1a). From Plumptre et al., (Plumptre et al. 2012), we extracted data for the 1986-2012 flowering anomalies in Lope, Gabon (Lat - 0.441156°, Long 11.524798°). Average monthly values were obtained from the same source for the proportion of trees flowering and the proportion of trees with ripe fruit in Goualougo, Nouabale Ndoki Park, Republic of the Congo (Lat 2.263195°, Long 16.591688°); the proportion of trees flowering; and the proportion of trees bearing ripe fruit in Lope, Gabon; the proportion of trees flowering and the proportion of trees bearing ripe fruit in Okapi National Park, Democratic Republic of the Congo (Lat 1.400931°, Long 28.577619°). We also obtained values for the annual proportion of trees fruiting in Kibale NP, Uganda 1970-2001 (Chapman et al. 2005). With an average monthly resolution, we extracted data for the percentage of trees with ripe fruit in Kibale National Park, Uganda between August 1990-September 2002 (Chapman et al. 2005). Arithmetic means were calculated for each year. From Philippon et al. (Philippon et al. 2007), we extracted data for the NDVI anomaly for West Africa (between Aug-Dec and between Jul-Dec 1982-2002). From Polansky and Boesch (Polansky and Boesch 2013), we also extracted the percentage of the forest community bearing fruit, and the count of number of species with peak fruiting times per month. Additional variables extracted (monthly, between 1994 and 2002, and annual averages) from graphs generated by Yamagiwa et al. (Yamagiwa et al. 2008), were a “fruit index” for fruit species consumed by Chimpanzees and Gorillas in both primary and secondary forest, in Kahuzi-Biega National Park, Democratic Republic of Congo (Lat. −1.963260°, Long 28.018609°). We further extracted data from the same publication, for the percentage of different plant species bearing fruit that contributed to the generation of the fruit indices (monthly data between 1998 and 2002, and annual arithmetic means). These include: *Allophylus* spp., *Bridelia bridelifolia*, *Cassipourea* spp., *Diospyros honleana*, *Ekebergia capensis*, *Ficus oreodryadum*, *Ficus thonningii*, *Maesa lanceolata*, *Myrianthus holstii*, *Newtonia buchananii*, *Psychotria palustris*, and *Syzygium parvifolium*.

All data are available as supplementary Appendix.

We analysed differences in climate and phenology variables between years without any spillover events, and years with at least one spillover event. The climate variables included were 10 yr average rainfall Kibale, avg. monthly max. temp Kibale -annual average-, DT90, DP01, DP05, DP10, CLDD, EMNT, EMXP, EMXT, MMNT, MMXT, MNTM, and TPCP. Phenology variables included were first and second independent components of NDVI (See Philippon et al. 2007 for details). First, we performed Kolmogorov-Smirnov tests on each quantitative variable to test for normality. For variables that were distributed normally, we subsequently performed univariate ANOVAs. For variables not normally distributed, we performed Kruskal-Wallis tests between years with and without spillover events. We subsequently box-cox transformed non-normally distributed variables. We then tested whether climate and phenology variables can predict the number of human and other animal Ebola spillover events. To this purpose, we performed a multiple regression in STATISTICA (Dell, Inc.) with number of spillover events as dependent variable, and climate and phenology variables as independent predictors. To further test whether phenology, or climate variables are better predictors for the recorded human + other animal spillover events, we generated and compared neural networks in Statistica (Statistica automated neural networks, SANN). First, principal components (PCs) were generated for 16 climate variables (Supplementary Table 1, 2), and for two phenology variables (Supplementary Table 3,4, for the time frame between 1970 and 2002). Subsequently, the data set was partitioned into a modelling dataset (years with even numbers) and a cross-validation data set (years with odd numbers). On the modelling dataset (70% training, 15% test, 15% validation data), we generated three models based on neural network time series regression, with different input variables, and the number of spillover events as target variable. First, a model was generated with climate + phenology PCs. Second, we generated a model using only phenology PCs. Third, we generated a model using only climate PCs as input. For model generation, we used the MLP (Multilayer Perceptron, Rumelhart et al. 1986) method with 500 networks and the 5 best networks retained. We applied weight decay to prevent overfitting and used 1 step for forecasting. For each analysis, the best model was chosen and subsequently deployed to the cross-validation dataset. We then compared the model estimates for the number of spillover events with the recorded number of spillover events in the cross-validation data set. Best-fit to the recorded number of spillover events was determined via correlations and t-tests for dependent samples. Monthly averages across years were available from a set of climate and phenology variables to determine seasonal patterns in the data. To reduce dimensionality in the data for analysis, we performed a principal component analysis. Thirteen climate variables were reduced to three PCs (Supplementary Tables 5, 6). 25 phenology variables were reduced to six principal components (Supplementary Tables 7,8). To explore the seasonal progression of these variables relative to one another, we plotted the number of spillovers per month, together with the average of climate PC values, and the average of phenology PC values. Between 1994 and 2002, monthly data was extracted for fruiting phenology, additionally to climate variables. For analysis, the monthly data partition was therefore cropped to only include monthly values from the years 1994-2002. To reduce dimensionality of climate and phenology variables, three principal component analyses were performed. Firstly, the eight climate variables were reduced to three principal components, that together explained 81.3% of the overall variance in this data set (See Supplementary Tables 9 and 10). From the set of four fruit index variables of Yamagiwa et al. (Yamagiwa et al. 2008), two principal components were extracted (Supplementary Tables 11 and 12). A principal component analysis for 12 single species variables recovered 5 principal components (Supplementary Tables 13 and 14). To test whether phenology and climate variables are predictive for Ebola spillover events in the monthly data partition, we performed a multiple regression.

## Results

Results of tests for normality for annual climate and phenology variables are shown in Table 1. Univariate ANOVAs (for normally distributed variables) and Kruskal-Wallis tests (for non-normally distributed variables, respectively, revealed significant differences in the climate variables 10 year avg. rainfall Kibale, DP01, DP05, DP10, EMNT, EMXP, MMNT, and TPCP between years with versus years without spillover events. Of the two phenology variables, IC2 NDVI significantly differed between the years with vs. without spillover events. Figure 1b shows box plots for these variables. Regression of climate and phenology variables against the number of human + other animal spillover events was overall significant (ANOVA results: R²= 0.3624, F(13,63)=2.7546, p<0.00374, Table 2). The residuals were distributed normally (not shown). Significant univariate predictors for this model were EMXT (p=0.049), and IC2 NDVI (p=0.012). To further determine the effectiveness of phenology versus climate variables in predicting spillover events, we generated and compared neural networks on principal components for the two types of predictor variables separately (phenology only, climate only), and combined (phenology and climate). Performance of each of the best models is shown in Supplementary Table 15. We do not aim for predictive quality of these models here - instead, we aimed to test for best fit between modelled data using different sets of input variables and observed data. For this purpose, we employed a paired t-test. When deploying each best model for the three sets of input variables, no model significantly differed from the observed values of Ebola spillover events, indicating an overall good fit (Figure 2a, results of t-test shown in Supplementary Table 16). The model with only phenology PCs as input variables performed best when deployed to the cross-validation data set, with the best correlation coefficient between predicted and observed spillover events (Figure 2b, r = 0.8860, p ≤ 0.000001).

**Table 1.** Significant differences in climate and phenology between years with no recorded spillover, and years with spillover of Ebola virus disease (to humans and animals). Shown are results of Kolmogorov-Smirnov tests (D-statistic, p) for variable normality, ANOVA (F-statistic, p, in the case of normality) or Kruskal-Wallis tests (H-Statistic, p in the case of non-normality). Variable type C - Climate, P - Phenology. For further variable abbreviations see Methods section.

| Variable type | Variable | K-S Test D | K-S Test p | ANOVA F | ANOVA p | KW-H | KW p | N Years |
| --- | --- | --- | --- | --- | --- | --- | --- | --- |
| C | 10 yr avg Rainfall Kibale | 0.1024 | n.s. | 10.0263 | **0.0025** |  |  | 60 |
| C | DP01 | 0.1595 | <0.1 |  |  | 5.4921 | **0.0191** | 59 |
| C | DP05 | 0.1405 | <0.2 |  |  | 8.8842 | **0.0029** | 59 |
| C | DP10 | 0.1335 | n.s. | 10.54 | **0.0020** |  |  | 59 |
| C | EMNT | 0.0693 | n.s. | 10.158 | **0.0024** |  |  | 55 |
| C | EMXP | 0.3016 | <0.01 |  |  | 10.3339 | **0.0013** | 55 |
| C | MMNT | 0.125 | n.s. | 6.1472 | **0.0163** |  |  | 57 |
| C | TPCP | 0.1972 | <0.05 |  |  | 19.911 | **0.00001** | 55 |
| P | IC2 NDVI anomaly(Jul-Dec) | 0.173 | n.s. | 5.3154 | **0.0326** |  |  | 21 |

**Table 2.** Regression of climate and phenology variables against annual spillover events in humans and animals. Variables were normalized with Box-cox transformation. The overall model is significant. ANOVA results: R²= 0.3624; F(13,63)=2.7546; p< 0.00374; Std.Error of estimate: 1.8584. Significant univariate predictors are shown in bold. Variable type C - Climate, P - Phenology. For further variable abbreviations see Methods section.

| Variable type | N=77 | b | Std.Err. of b | t(63) | p-value |
| --- | --- | --- | --- | --- | --- |
| C | CLDD | -0.0009 | 0.0006 | -1.4456 | 0.1533 |
| C | EMNT | 0.0805 | 0.1675 | 1.1183 | 0.2677 |
| C | EMXP | 1.6249 | 6.7085 | 0.2422 | 0.8094 |
| **C** | **EMXT** | **0.1459** | **0.0728** | **2.0053** | **0.0492** |
| C | MMNT | -0.0294 | 0.1256 | -0.2340 | 0.8157 |
| C | MMXT | -0.2098 | 0.1472 | -1.4258 | 0.1589 |
| C | MNTM | 0.0702 | 0.1874 | 0.3749 | 0.7090 |
| C | TPCP | -0.0557 | 0.1473 | -0.3781 | 0.7066 |
| P | %of trees with ripe fruit Kibale | 0.4504 | 0.3796 | 1.1865 | 0.2399 |
| P | Proportion of population fruiting in Kibale | -0.0169 | 0.0557 | -0.3039 | 0.7622 |
| P | NDVI anomaly (Aug-Nov) | -0.5641 | 0.5007 | -1.1265 | 0.2642 |
| **P** | **NDVI anomaly (Jul-Dec)** | **1.4756** | **0.5691** | **2.5931** | **0.0118** |
| P | Flowering Anomalies Lope Gabon | 0.1351 | 0.0818 | 1.6509 | 0.1037 |

**Figure 2.**
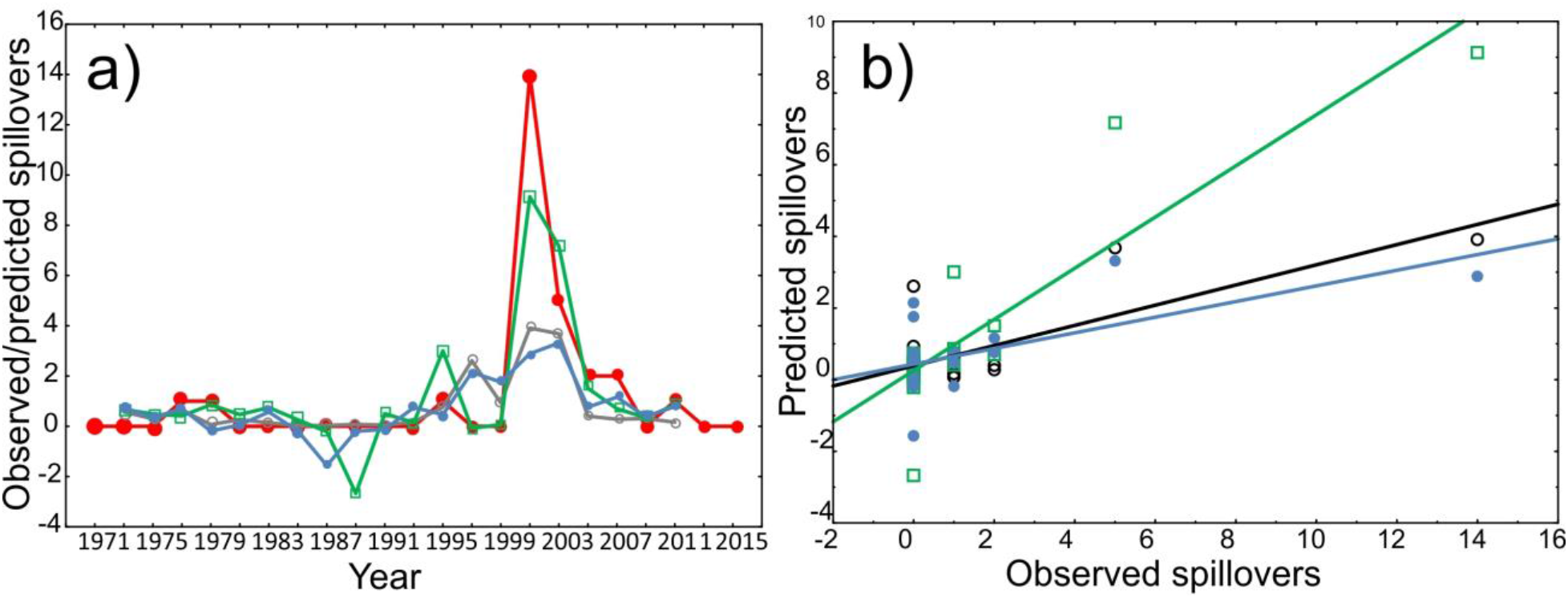
Performance for annual time series regression of animal/human spillover events from Neural Network modeling (SANN) using only climate PCs, only phenology PCs, and climate and phenology PCs combined as sets of input variables. Panel a) shows time series predictions for the observed number of human and animal spillover (closed large circles/red), vs. the predicted numbers from best retained models (500 network iterations) of climate variables only (small closed circles/blue), phenology variables only (open squares/green), and climate/phenology combined as input data sets (open circles/grey). Panel b) shows correlation scatter plots between observed spillovers and predicted spillovers using models with inputs: climate variables only (small closed circles/blue), phenology variables only (open squares/green), and climate/phenology combined as input data sets (open circles/grey). Correlation with Climate + Phenology model r=0.7663, p=0.00008; Correlation with only climate model r = 0.6254, p=0.0032; Correlation with only phenology r = 0.8860, p≤0.000001.

Using average values of phenology and of climate principal components, line plots for seasonality (Figure 3) show a (previously reported) pattern of seasonality of Ebola spillover events. Our strategy to record both human and other animal spillover events, as well as recording independent epidemic chains as separate spillover events, reveals a clear seasonal dynamic of recorded spillovers, phenology, and climate variables (Figure 3).

**Figure 3.**
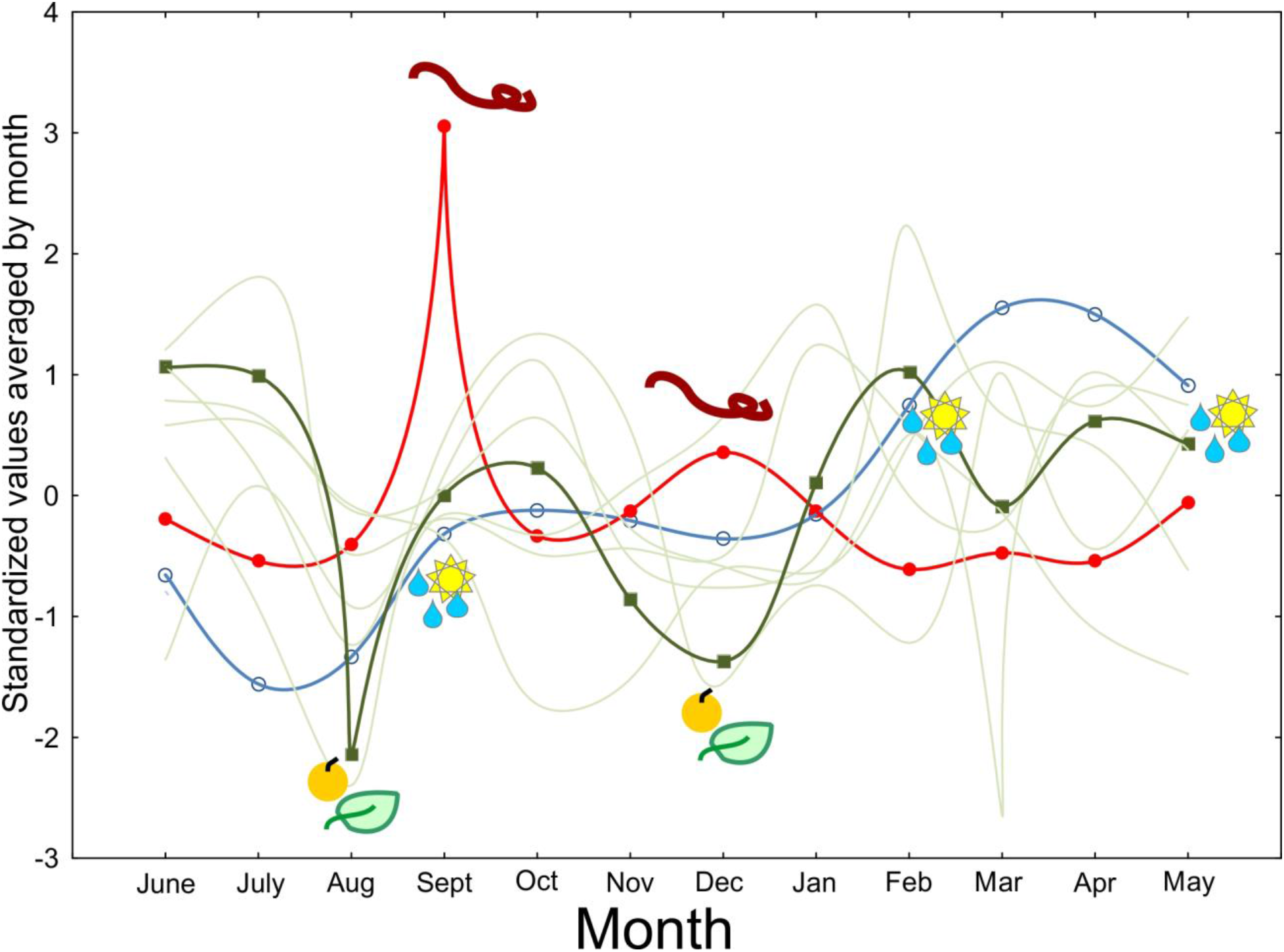
Seasonality of Ebola virus spillover events (humans + animals, standardized values, closed circles), the standardized average of climate PCs 1-3 (open circles) and the standardized average of phenology PCs 1-6 (closed squares). Standardized values for single phenology PCs 1-6 are shown as background lines (no symbols). Both September and December spikes in recorded human + animal Ebola virus spillover events (virus symbols) are associated with lower values of the average of phenology PCs 1-6 (fruit/leaf symbols), but only for the September spike with average climate variable changes (weather symbols). Curves are spline-fitted.

The Yamagiwa et al. (Yamagiwa et al. 2008) data set allowed a detailed investigation into which kinds of plants are contributing to the inter-annual and seasonal variation in phenology that we found to be associated with Ebola spillover events. Multiple regression revealed that for the investigated time period (which includes a total of 57 spillover events), climate variation does not significantly predict the number of monthly spillovers. While the “fruit index” variables also were not significant predictors, three of the five plant species PCs significantly predicted spillover events (Table 3). The overall regression model was significant (ANOVA for overall goodness of fit: R²= 0.2368, F (11,93)=2.6238, p<0.00583, Std.Error of estimate, 0.9740). Variables with significant factor loadings for these three PCs were the proportion of trees bearing fruit: *Ekebergia capensis, Ficus oreodryadum, Ficus thonningii*, and *Newtonia buchananii* (Supplementary Table 13). A comparison of means and errors for the proportion of these plants fruiting for the analysed time series 1993-2002 is shown in Figure 4.

**Table 3.** Multiple regression analysis for human + animal spillover events against climate, fruit, and plant variables (Monthly data partition). The overall model was significant (ANOVA for overall goodness of fit: R²= 0.2368; F(11,93)=2.6238; p < 0.00583; Std.Error of estimate: 0.974). Variable type C - Climate, P -Phenology. For further variable abbreviations see Methods section.

| Variable type | Variable | b | Std.Err. of b | t(93) | p-value |
| --- | --- | --- | --- | --- | --- |
| C | M_Climate PC1 | -0.0483 | 0.1058 | -0.4562 | 0.6493 |
| C | M_Climate PC2 | -0.1165 | 0.0972 | -1.1983 | 0.2338 |
| C | M_Climate PC3 | 0.0767 | 0.1009 | 0.7595 | 0.4495 |
| P | M_Fruit PC1 | -0.0703 | 0.1224 | -0.5747 | 0.5669 |
| P | M_Fruit PC2 | 0.1387 | 0.2136 | 0.6495 | 0.5176 |
| P | M_Plant PC1 | -0.1622 | 0.1025 | -1.5822 | 0.1170 |
| **P** | **M_Plant PC2** | **0.2813** | **0.1106** | **2.5428** | **0.0126** |
| **P** | **M_Plant PC3** | **-0.2731** | **0.1010** | **-2.7050** | **0.0081** |
| **P** | **M_Plant PC4** | **-0.2760** | **0.1111** | **-2.4834** | **0.0148** |
| P | M_Plant PC5 | -0.0211 | 0.0981 | -0.2154 | 0.8299 |

**Figure 4.**
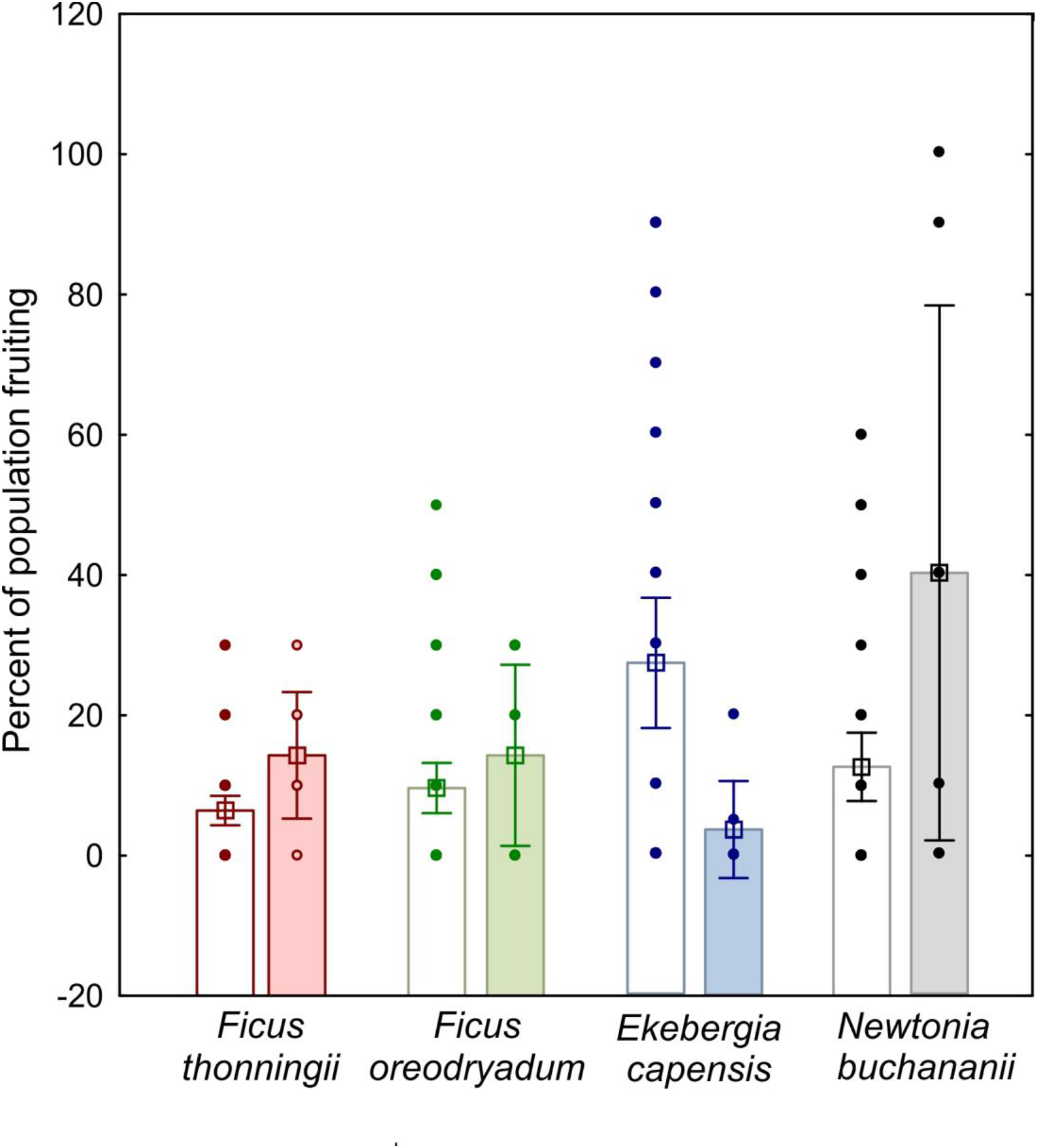
Changes in the proportion of plants bearing fruits between months with no recorded spillovers (open bars) and months with recorded spillovers (closed bars). Dots represent raw data points, column height means, bars represent errors. Plant species shown here were significant predictors in the regression model for number of spillovers per month for the period 1993-2002 in Kahuzi-Biega National Park. Original data from Yamagiwa et al. (2008).

## Discussion

We found significant differences in climate and vegetation parameters between years with and years without Ebola virus spillover events. Years with spillover events were characterized by generally drier conditions (DP01, DP05, EMXP, TPCP), as well as higher minimum temperatures (EMNT, MMNT). In contrast, the anomaly of the vegetation index in the months from July - December was higher, combined with a higher average of rainfall (measured in Kibale, Uganda). An overall model of climate and phenology variables as predictors for the number of human + other animal spillovers in a given year was significant. EMXT, the highest monthly daily maximum temperature, was a significant predictor for the number of Ebola spillover events in humans and animals. These results align to previous studies that have found a climatic dimension for Ebola spillover events (Tucker et al. 2002; Pigott et al. 2014; Schmidt et al. 2017). In our study, however, we could additionally show that plant phenology variables (including the anomaly of Normalized Difference Vegetation Index (NDVI) between July and December, the proportion of population fruiting in Kibale NP, Uganda, and the flowering anomalies in Lope, Gabon), informed neural network models with a superior fit to the data than when climate or climate in conjunction with phenology variables were used as inputs. Previous studies using Normalized Difference Vegetation Index or Enhanced Vegetation Index have not yet considered their predictive power for Ebola spillover events independently from climate variables (see for example Schmidt et al. 2017).

Aligned to previous reports (Alexander et al. 2015), we found a pronounced seasonality in recorded human and other animal spillover events. This seasonal association between Ebola spillover events and a multitude of phenology variables describing seasonality of flowering and fruiting was not previously reported, and corroborates the newly found close association between inter-annual Ebola spillover events and interannual variation in vegetation and phenology variables. The seasonality in spillover events is thought to be associated with transitions between wet-to-dry or dry-to-wet conditions (Tucker et al. 2002; Pinzon et al. 2004; Groseth et al. 2007; Altizer et al. 2013; Schmidt et al. 2017), and thus may be associated with spatiotemporal fluctuations of the African Monsoon (Cornforth 2013). While the September spike in human + other animal spillover events was associated with such a transition of climatic variables, the December spike was not. Other time points with shifts in climatic conditions were vice versa not associated with increases in Ebola spillover events. In contrast, both spikes in Ebola spillovers are associated with lower values in plant phenology variables, lending additional support to the multi-emergence hypothesis. Previous studies have found seasonal and yearly oscillations of fruiting status within specific localities, with a trend that is generated by differences in peak fruiting times among individual species (Anderson et al. 2005; Polansky and Boesch 2013). Most species fruit only for one month per year (Anderson et al. 2005). Fruit abundance of species within the Tai forest (Ivory Coast) plant community has been found to increase over the past decade, with no apparent correlation with local climatic patterns (Polansky and Boesch 2013). Basabose (Basabose 2002) also reported significant differences in the mean percentage of trees fruiting at Kahuzi-Biega National Park between 1994 and 2000, both between the dry and the wet season, as well as between years. Data from both these localities were part of our analysis. Local flowering and fruiting patterns consequently might reflect components of the local environment of the Ebola virus reservoir that differ from climatic variables, represented by climate layers.

Without in-depth knowledge of the proximate mechanism of Ebola spillovers, we could here show a covariation between ecological variables related to food resources, and Ebola spillovers to animal hosts. For example, “virus shedding” (McFarlane et al. 2011; Plowright et al. 2015), the process of bats transmitting virus particles to plant matter which is then uptaken by other animals, provides a possible functional link between phenology and virus spillover. Plant phenology variables that significantly predicted monthly spill-over events included *Ficus spp.* fruit, leaves, and stems which are consumed by Gorilla and Chimpanzee. *Ekebergia capensis* fruit and *Newtonia buchananii* seed are likewise consumed by Gorilla and Chimpanzee in Kahuzi-Biega National Park (Democratic Republic of Congo, (Yamagiwa and Basabose 2006). *Ficus thonningii* constitutes an important food source for African frugivores (Bleher et al. 2003; Kirika et al. 2008). Seasonally dry climate has shown to cause synchronicity for flowering, but not fruiting in this species (St J. Damstra et al. 1996). It has been proposed as a keystone species for frugivores, as it bears fruit during periods of fruit scarcity (Bleher et al. 2003). We could here show an association between Ebola spillovers and phenology of plants that constitute a seasonally varying food source for susceptible species. Plant species like *F. thonningii* could serve as preliminary focal species to test the hypothesis that competition for fruit or other plant matter in dry periods increase the likelihood of spillover events. Our results on the link between Ebola spillovers and forest phenology is mirrored by results from another viral haemorrhagic fever in Belgium; there, dynamics of vegetation phenology and their alteration under climate change have been found to influence the dynamics of Puumala virus (causing the disease Nephropathia endemica, (Barrios et al. 2010). In conclusion, both Barrios et al. (2010) and our present study support the multi-emergence hypothesis for virus spillover events. The phenology source data analysed in this study, however, were far from comprehensive. Clearly, to improve the understanding of the importance of phenology for predicting disease emergence and the quality of predictive models, more data are needed that spatially match locations of Ebola virus emergence. Phenology can vary locally, and seasonally (Bleher et al. 2003), so that this information cannot be replaced by remote-sensing data. We therefore highlight the importance of ecological field work additional to remote sensing in providing important data on phenology variables related to the ecosystem ecology of the unknown natural reservoir of Ebola virus.

## Conclusion

Data on local fruiting patterns, collected locally, could constitute a useful indicator for the likelihood of impending Ebola spillover events and could readily be classified through cost-effective transect surveys. The most recent Ebola outbreak in May 2017 in Likati (Bas-Uele Province, Democratic Republic of Congo) suggests that the emergence of Ebola virus is a recurring event that necessitates the generation of upstream models for disease management that is sensitive to economic cost and cost of human lives (Schar and Daszak 2014). We therefore encourage researchers to make unpublished African phenology data available to the public domain to aid in understanding this dimension of ecological and environmental variables related to spillover events of emerging infectious diseases.

## Acknowledgements

A 2015 Mathematics-Biology Data Analytics Workshop at Bethune-Cookman University provided the venue for initial discussion, planning and data generation for this project. We thank all student and faculty participants for valuable discussions. The Mathematics-Biology Data Analytics Workshop was funded by National Science Foundation project HBCU-UP Targeted Infusion project HRD-1435186.

## References

Adole T, Dash J, Atkinson PM (2016) A systematic review of vegetation phenology in Africa. Ecol Inform 34:117–128.

Alexander KA, Sanderson CE, Marathe M, Lewis BL, Rivers CM, Shaman J, Drake JM, Lofgren E, Dato VM, Eisenberg MC, Eubank S (2015) What factors might have led to the emergence of Ebola in West Africa? PLoS Negl Trop Dis 9:e0003652.

Altizer S, Ostfeld RS, Johnson PTJ, Kutz S, Harvell CD (2013) Climate change and infectious diseases: from evidence to a predictive framework. Science 341:514–519.

Amblard J, Obiang P, Edzang S, Prehaud C, Bouloy M, Guenno BL (1997) Identification of the Ebola virus in Gabon in 1994. Lancet 349:181–182.

Anderson DP, Nordheim E, Moermond TC, Bi G, Zoro B, Boesch C, Others (2005) Factors influencing tree phenology in taï national park, Côte d’Ivoire. Biotropica 37:631–640.

Baize S, Pannetier D, Oestereich L, Rieger T, Koivogui L, Magassouba N’faly, Soropogui B, Sow MS, Keïta S, De Clerck H, Tiffany A, Dominguez G, Loua M, Traoré A, Kolié M, Malano ER, Heleze E, Bocquin A, Mély S, Raoul H, Caro V, Cadar D, Gabriel M, Pahlmann M, Tappe D, Schmidt-Chanasit J, Impouma B, Diallo AK, Formenty P, Van Herp M, Günther S (2014) Emergence of Zaire Ebola virus disease in Guinea. N Engl J Med 371:1418–1425.

Bärbel Bleher, Potgieter CJ, C. J. Potgeiter, Johnson DN, Böhning-Gaese K (2003) The Importance of Figs for Frugivores in a South African Coastal Forest. J Trop Ecol 19:375–386.

Barrios JM, Verstraeten WW, Maes P, Clement J, Aerts J-M, Haredasht SA, Wambacq J, Lagrou K, Ducoffre G, Van Ranst M, Berckmans D, Coppin P (2010) Satellite derived forest phenology and its relation with nephropathia epidemica in Belgium. Int J Environ Res Public Health 7:2486–2500.

Basabose AK (2002) Diet composition of chimpanzees inhabiting the montane forest of Kahuzi, Democratic Republic of Congo. Am J Primatol 58:1–21.

Bausch DG, Schwarz L (2014) Outbreak of ebola virus disease in Guinea: where ecology meets economy. PLoS Negl Trop Dis 8:e3056.

Boyd E, Cornforth RJ, Lamb PJ, Tarhule A, Issa Lélé M, Brouder A (2013) Building resilience to face recurring environmental crisis in African Sahel. Nat Clim Chang 3:631–637.

Chapman CA, Chapman LJ, Struhsaker TT, Zanne AE, Clark CJ, Poulsen JR (2005) A Long-Term Evaluation of Fruiting Phenology: Importance of Climate Change. J Trop Ecol 21:31–45.

Cornforth RJ (2013) West African Monsoon 2012. Weather 68:256–263.

Fisher JB, Sikka M, Sitch S, Ciais P, Poulter B, Galbraith D, Lee J-E, Huntingford C, Viovy N, Zeng N, Ahlström A, Lomas MR, Levy PE, Frankenberg C, Saatchi S, Malhi Y (2013) African tropical rainforest net carbon dioxide fluxes in the twentieth century. Philos Trans R Soc Lond B Biol Sci 368:20120376.

Formenty P, Hatz C, Le Guenno B, Stoll A (1999) Human infection due to Ebola virus, subtype Cote d’Ivoire: clinical and biologic presentation.

Georges AJ, Leroy EM, Renaut AA, Benissan CT, Nabias RJ, Ngoc MT, Obiang PI, Lepage JP, Bertherat EJ, Bénoni DD, Wickings EJ, Amblard JP, Lansoud-Soukate JM, Milleliri JM, Baize S, Georges-Courbot MC (1999) Ebola hemorrhagic fever outbreaks in Gabon, 1994-1997: epidemiologic and health control issues. J Infect Dis 179 Suppl 1:S65–75.

Groseth A, Feldmann H, Strong JE (2007) The ecology of Ebola virus. Trends Microbiol 15:408–416.

Heymann DL, Weisfeld JS, Webb PA, Johnson KM, Cairns T, Berquist H (1980) Ebola hemorrhagic fever: Tandala, Zaire, 1977–1978. J Infect Dis 142:372–376.

International Commission (1978) Ebola haemorrhagic fever in Zaire, 1976. Bull World Health Organ 56:271–293.

Khan AS, Tshioko FK, Heymann DL, Le Guenno B, Nabeth P, Kerstiëns B, Fleerackers Y, Kilmarx PH, Rodier GR, Nkuku O, Rollin PE, Sanchez A, Zaki SR, Swanepoel R, Tomori O, Nichol ST, Peters CJ, Muyembe-Tamfum JJ, Ksiazek TG (1999) The reemergence of Ebola hemorrhagic fever, Democratic Republic of the Congo, 1995. Commission de Lutte contre les Epidémies à Kikwit. J Infect Dis 179 Suppl 1:S76–86.

Kim St J. Damstra, Richardson S, Reeler B (1996) Synchronized Fruiting Between Trees of Ficus thonningii in Seasonally Dry Habitats. J Biogeogr 23:495–500.

Kirika JM, Bleher B, Böhning-Gaese K, Chira R, Farwig N (2008) Fragmentation and local disturbance of forests reduce frugivore diversity and fruit removal in Ficus thonningii trees. Basic Appl Ecol 9:663–672.

Lahm SA, Kombila M, Swanepoel R, Barnes RFW (2007) Morbidity and mortality of wild animals in relation to outbreaks of Ebola haemorrhagic fever in Gabon, 1994–2003. Trans R Soc Trop Med Hyg 101:64–78.

Lamunu M, Lutwama JJ, Kamugisha J, Opio A, Nambooze J, Ndayimirije N, Okware S (2004) Containing a haemorrhagic fever epidemic: the Ebola experience in Uganda (October 2000-January 2001). Int J Infect Dis 8:27–37.

Leendertz SAJ, Gogarten JF, Düx A, Calvignac-Spencer S, Leendertz FH (2016) Assessing the Evidence Supporting Fruit Bats as the Primary Reservoirs for Ebola Viruses. EcoHealth 13:18–25.

Le Guenno B, Formenty P, Formentry P, Wyers M, Gounon P, Walker F, Boesch C (1995) Isolation and partial characterisation of a new strain of Ebola virus. Lancet 345:1271–1274.

Leroy EM, Epelboin A, Mondonge V, Pourrut X, Gonzalez J-P, Muyembe-Tamfum J-J, Formenty P (2009) Human Ebola outbreak resulting from direct exposure to fruit bats in Luebo, Democratic Republic of Congo, 2007. Vector Borne Zoonotic Dis 9:723–728.

Leroy EM, Rouquet P, Souquiere S, Kilbourne A, Bermejo M, Smit S (2004) Multiple Ebola Virus Transmission Events and Rapid Decline of Central African Wildlife. Science 303:387–390.

Leroy E, Gonzalez J-P (2012) Filovirus research in Gabon and Equatorial Africa: The experience of a research center in the heart of Africa. Viruses 4:1592–1604.

MacNeil A, Farnon EC, Wamala J, Okware S, Cannon DL, Reed Z, Towner JS, Tappero JW, Lutwama J, Downing R, Nichol ST, Ksiazek TG, Rollin PE (2010) Proportion of deaths and clinical features in Bundibugyo Ebola virus infection, Uganda. Emerg Infect Dis 16:1969–1972.

McFarlane R, Becker N, Field H (2011) Investigation of the climatic and environmental context of Hendra virus spillover events 1994–2010. PLoS One 6:e28374.

Milleliri JM, Tévi-Benissan C, Baize S, Leroy E, Georges-Courbot MC (2004) [Epidemics of Ebola haemorrhagic fever in Gabon (1994-2002). Epidemiologic aspects and considerations on control measures]. Bull Soc Pathol Exot 97:199–205.

Muyembe T, Kipasa M (1995) Ebola haemorrhagic fever in Kikwit, Zaire. International Scientific and Technical Committee and WHO Collaborating Centre for Haemorrhagic Fevers. Lancet 345:1448.

Nkoghe D, Kone ML, Yada A, Leroy E (2011) A limited outbreak of Ebola haemorrhagic fever in Etoumbi, Republic of Congo, 2005. Trans R Soc Trop Med Hyg 105:466–472.

Okware SI, Omaswa FG, Zaramba S, Opio A, Lutwama JJ, Kamugisha J, Rwaguma EB, Kagwa P, Lamunu M (2002) An outbreak of Ebola in Uganda. Trop Med Int Health 7:1068–1075.

Olival KJ, Hosseini PR, Zambrana-Torrelio C, Ross N, Bogich TL, Daszak P (2017) Host and viral traits predict zoonotic spillover from mammals. Nature 546:646–650.

Onyango CO, Opoka ML, Ksiazek TG, Formenty P, Ahmed A, Tukei PM, Sang RC, Ofula VO, Konongoi SL, Coldren RL, Grein T, Legros D, Bell M, De Cock KM, Bellini WJ, Towner JS, Nichol ST, Rollin PE (2007) Laboratory diagnosis of Ebola hemorrhagic fever during an outbreak in Yambio, Sudan, 2004. J Infect Dis 196 Suppl 2:S193–8.

Peterson AT, Bauer JT, Mills JN (2004) Ecologic and geographic distribution of filovirus disease. Emerg Infect Dis 10:40–47.

Philippon N, Jarlan L, Martiny N, Camberlin P, Mougin E (2007) Characterization of the Interannual and Intraseasonal Variability of West African Vegetation between 1982 and 2002 by Means of NOAA AVHRR NDVI Data. J Clim 20:1202–1218.

Pigott DM, Golding N, Mylne A, Huang Z, Henry AJ, Weiss DJ, Brady OJ, Kraemer MUG, Smith DL, Moyes CL, Bhatt S, Gething PW, Horby PW, Bogoch II, Brownstein JS, Mekaru SR, Tatem AJ, Khan K, Hay SI (2014) Mapping the zoonotic niche of Ebola virus disease in Africa. Elife 3:e04395.

Pinzon JE, Wilson JM, Tucker CJ, Arthur R, Jahrling PB, Formenty P (2004) Trigger events: enviroclimatic coupling of Ebola hemorrhagic fever outbreaks. Am J Trop Med Hyg 71:664–674.

Plowright RK, Eby P, Hudson PJ, Smith IL, Westcott D, Bryden WL, Middleton D, Reid PA, McFarlane RA, Martin G, Tabor GM, Skerratt LF, Anderson DL, Crameri G, Quammen D, Jordan D, Freeman P, Wang L-F, Epstein JH, Marsh GA, Kung NY, McCallum H (2015) Ecological dynamics of emerging bat virus spillover. Proc Biol Sci 282:20142124.

Plumptre AJ, Shoo L, Polansky L, Strindberg S, Boesch K, Breuer T, Bujo F, Chapman CA, Hart MCT, Hockemba M, Hosaka K, Jeffrey K, Ndangalasi NIH, Ndolo S, O’Brien T, Pusey A, Ssali. F (2012) Changes in Tree Phenology across Africa: A comparison across 17 sites. WCS Albertine Rift program

Polansky L, Boesch C (2013) Long-term Changes in Fruit Phenology in a West African Lowland Tropical Rain Forest are Not Explained by Rainfall. Biotropica 45:434–440.

Pourrut X, Kumulungui B, Wittmann T, Moussavou G, Délicat A, Yaba P, Nkoghe D, Gonzalez J-P, Leroy EM (2005) The natural history of Ebola virus in Africa. Microbes Infect 7:1005–1014.

Rohatgi A (2016) WebPlotDigitizer 3.10. Accessible at http://arohatgi.info/WebPlotDigitizer.

Rouquet P, Froment J-M, Bermejo M, Kilbourn A, Karesh W, Reed P, Kumulungui B, Yaba P, Délicat A, Rollin PE, Leroy EM (2005) Wild animal mortality monitoring and human Ebola outbreaks, Gabon and Republic of Congo, 2001–2003. Emerg Infect Dis 11:283–290.

Rumelhart DE, Hinton GE, Williams RJ (1986) Parallel Distributed Processing: Explorations in the Microstructure of Cognition, Vol. 1. In: Rumelhart DE, McClelland JL, PDP Research Group C (eds). MIT Press, Cambridge, MA, USA, pp 318–362.

Schar D, Daszak P (2014) Ebola economics: the case for an upstream approach to disease emergence. Ecohealth 11:451–452.

Schmidt JP, Park AW, Kramer AM, Han BA, Alexander LW, Drake JM (2017) Spatiotemporal Fluctuations and Triggers of Ebola Virus Spillover. Emerg Infect Dis 23:415–422.

Shoemaker T, MacNeil A, Balinandi S, Campbell S, Wamala JF, McMullan LK, Downing R, Lutwama J, Mbidde E, Ströher U, Rollin PE, Nichol ST (2012) Reemerging Sudan Ebola virus disease in Uganda, 2011. Emerg Infect Dis 18:1480–1483.

Taylor DJ, Leach RW, Bruenn J (2010) Filoviruses are ancient and integrated into mammalian genomes. BMC Evol Biol 10:193.

Towner JS, Sealy TK, Khristova ML, Albariño CG, Conlan S, Reeder SA, Quan P-L, Lipkin WI, Downing R, Tappero JW, Okware S, Lutwama J, Bakamutumaho B, Kayiwa J, Comer JA, Rollin PE, Ksiazek TG, Nichol ST (2008) Newly discovered ebola virus associated with hemorrhagic fever outbreak in Uganda. PLoS Pathog 4:e1000212.

Tucker CJ, Wilson JM, Mahoney R, Anyamba A, Linthicum K, Myers MF (2002) Climatic and ecological context of the 1994–1996 Ebola outbreaks. Photogramm Eng Remote Sens 68:147–152.

Wamala JF, Lukwago L, Malimbo M, Nguku P, Yoti Z, Musenero M, Amone J, Mbabazi W, Nanyunja M, Zaramba S, Opio A, Lutwama JJ, Talisuna AO, Okware SI (2010) Ebola hemorrhagic fever associated with novel virus strain, Uganda, 2007–2008. Emerg Infect Dis 16:1087–1092.

Weber DS, Alroy KA, Scheiner SM (2017) Phylogenetic Insight into Zika and Emerging Viruses for a Perspective on Potential Hosts. Ecohealth 14:214–218.

Wittmann TJ, Biek R, Hassanin A, Rouquet P, Reed P, Yaba P, Pourrut X, Real LA, Gonzalez J-P, Leroy EM (2007) Isolates of Zaire ebolavirus from wild apes reveal genetic lineage and recombinants. Proc Natl Acad Sci U S A 104:17123–17127.

Yamagiwa J, Basabose AK (2006) Diet and seasonal changes in sympatric gorillas and chimpanzees at Kahuzi–Biega National Park. Primates 47:74–90.

Yamagiwa J, Basabose AK, Kaleme KP, Yumoto T (2008) Phenology of fruits consumed by a sympatric population of gorillas and chimpanzees in Kahuzibiega National Park, Democratic Republic of Congo. African Study Monographs 3–22.

